# Human alveolar lining fluid from the elderly promotes *Mycobacterium tuberculosis* growth in alveolar epithelial cells and bacterial translocation into the cytosol

**DOI:** 10.1101/2021.05.12.443884

**Authors:** Angélica M. Olmo-Fontánez, Julia M. Scordo, Andreu Garcia-Vilanova, Diego Jose Maselli, Jay I. Peters, Blanca I. Restrepo, Daniel L. Clemens, Joanne Turner, Larry S. Schlesinger, Jordi B. Torrelles

**Affiliations:** Population Health and Host Pathogen Interactions Programs, Texas Biomedical Research Institute, San Antonio, TX, 78227, USA; Integrated Biomedical Sciences Program, University of Texas Health Science Center at San Antonio, TX, 78229, USA; Sam and Ann Barshop Institute for Longevity and Aging Studies, University of Texas Health Science Center at San Antonio, TX, 78229, USA; Division of Pulmonary and Critical Care Medicine, School of Medicine, University of Texas Health Science Center at San Antonio, TX, 78229, USA; University of Texas Health Science Center at Houston, School of Public Health, Brownsville campus, Brownsville, TX 78520, USA; and South Texas Diabetes and Obesity Institute, University of Texas Rio Grande Valley, Edinburg, TX 78541, USA; University of California, Los Angeles Health Sciences, Los Angeles, CA, 90095, USA

**Keywords:** *Mycobacterium tuberculosis*, aging, alveolar epithelial cells, alveolar lining fluid, cytosol, tuberculosis

## Abstract

The elderly population is at significant risk of developing respiratory diseases, including tuberculosis (TB) caused by the airborne *Mycobacterium tuberculosis* (*M.tb*). Once *M.tb* reaches the alveolar space, it contacts alveolar lining fluid (ALF) which dictates host cell interactions. We previously determined that age-associated dysfunctionality in human ALF soluble innate components lead to accelerated *M.tb* growth within human alveolar macrophages. Here we determined the impact of human ALF on *M.tb* infection of alveolar epithelial cells (ATs), another critical cellular determinant of infection. We observed that E-ALF-exposed *M.tb* had significantly increased intracellular growth in ATs compared to adult ALF (A-ALF)-exposed bacteria. Despite this, there were no alterations in AT inflammatory mediators or cell activation. However, exposure to E-ALF altered endosomal trafficking of *M.tb*, driving bacterial translocation to both endosomal and cytosolic compartments in ATs. Our results indicate that exposure of *M.tb* to E-ALF promotes translocation of bacteria into the AT cytosol as a potential favorable niche for rapid bacterial growth and at the same time dampens AT’s immune responses. Thus, our findings highlight the influence of the elderly lung mucosa on *M.tb* infection of ATs, an unexplored contributing factor to the elderly population’s increased susceptibility of developing active TB disease.

## INTRODUCTION

Tuberculosis (TB) is one of the leading causes of death due to an infectious disease and is considered a global threat killing over 4,500 people every day.^1^ The risk of TB susceptibility and mortality is significantly increased in individuals aged 65 and older.^2, 3^ TB is caused by airborne *Mycobacterium tuberculosis* (*M.tb*), transmitted primarily by inhalation, where it is deposited into the distal portion of the airways and alveoli. In this environment, *M.tb* encounters the lung mucosa, or alveolar lining fluid (ALF), which contains soluble innate factors such as surfactant proteins A and D (SP-A/SP-D), hydrolytic enzymes, complement, lipids, and others, which activate subsequent innate and adaptive immune responses.^4-6^

As we age, changes to soluble components of the innate immune system in ALF may contribute to the increased susceptibility of the elderly population to TB.^7, 8^ Published studies from our labs found that ALF from elderly humans and old mice have increased levels of pro-inflammatory and pro-oxidative mediators, impacting *M.tb* infection outcomes *in vitro* and *in vivo*.^9^ Human macrophages infected with *M.tb* exposed to elderly human ALF (E-ALF) have reduced control of infection and altered intracellular trafficking with fewer phagosome–lysosome fusion events. These observations were reversed when E-ALF was replenished with functional SP-A/SP-D, supporting the importance of the functionality of ALF innate components in *M.tb* control.^9^ This was also observed *in vivo*, where *M.tb* that had been exposed to E-ALF grew faster and induced more lung immunopathology in young infected mice.^9^

Most studies focus on the role of ALF in altering *M.tb*-phagocyte interactions; ^4, 10, 11^ however, it is critical to understand the impact of ALF on *M.tb* infection of non-professional phagocytes, in particular, alveolar epithelial type cells (ATs).^12, 13^ ATs are the most prevalent cell population that covers the internal surface area of the alveolar environment.^14^ The alveolar epithelium contains two main epithelial cell types that maintain alveolus integrity, preventing microbial dissemination. Type I ATs are the most abundant cell type, providing a structural role in shaping the alveolus and allowing for gas exchange.^14^Type II ATs are spherical pneumocytes that comprise less than 5% of the surface area yet constitute 60% of the ATs and play an essential role in host defense and in maintaining alveolar homeostasis by secreting and recycling ALF components such as surfactant proteins, hydrolases, and mucosal antibodies, among others.^15-17^

When *M.tb* reaches the alveoli it first interacts with ATs and alveolar macrophages, with subsequent invasion and replication within the alveolar epithelial barrier.^18^ *M.tb* expresses a variety of virulence factors such as a heparin-binding hemagglutinin (HBHA) and malate synthase that promote adherence and entry into ATs.^19, 20^ Given that ATs are non-professional phagocytes, they are proposed to provide a protective niche that enables *M.tb* to replicate and elude an innate immune response. Nonetheless, ATs also participate in immune responses involved in controlling *M.tb* infection by producing pro-inflammatory cytokines (TNF, IL-8, and GM-CSF), thereby potentiating cellular crosstalk and activation of alveolar macrophages leading to an increase in their antimycobacterial activity.^21^ An additional host defense mechanism of ATs is the secretion of innate immune molecules, *e.g*., SP-A, SP-D, complement component 3 (C3), antimicrobial peptides, antibodies, and hydrolases, among others into the ALF that exhibits an essential role facilitating cell recruitment, microbial killing^16^ and even driving the differential outcome of *M.tb* infection in ATs.^13^ We recently found that *M.tb* exposed to ALF from healthy adults vary in growth rates within ATs, which was dependent on ALF protein oxidation levels and function.^13^ Based on these findings and our characterization of the elderly lung environment, we aimed to determine the impact of the elderly lung mucosa on *M.tb* infection of ATs. We provide evidence that *M.tb* exposure to E-ALF drives increased *M.tb* replication and growth in ATs, as well as *M.tb* translocation to both endosomal and cytosolic compartments in ATs, keeping unaltered AT cell death and early immune responses against the infection. These findings indicate that E-ALF promotes *M.tb* growth within ATs potentially by exploiting the AT cytosol as a protective replicative niche for *M.tb*.

## RESULTS

### *M.tb* exposure to elderly human ALF drives increased bacterial intracellular growth in alveolar epithelial cells (ATs) *in vitro*

Our prior studies have shown that *M.tb* exposure to ALF from elderly individuals (E-ALF) accelerates the growth of *M.tb* within human alveolar macrophages and human monocyte-derived macrophages.^9^ Here we observed that *M.tb* previously exposed to E-ALF also demonstrates significantly increased intracellular growth in ATs when compared to A-ALF-exposed *M.tb* (**Fig. 1A**). This increased bacterial growth of E-ALF-exposed *M.tb* was not due to differences in the inoculum used and/or uptake by ATs, because the inoculum of *M.tb* exposed to E-ALF or A-ALF and levels of uptake were equivalent (**Fig. 1B-C**). We further confirmed that intracellular growth differences within ATs were not due to changes in AT cell viability after infection with *M.tb* exposed to either A-ALF or E-ALF over the infection period (120 h) (**Suppl. Fig. S1**). We next explored the surface expression of e-Cadherin (CD324) on ATs in the different treatment groups. Our results showed high e-Cadherin expression throughout the infection, indicating that cell-cell interfaces between ATs were stable,^22^ consistent with high cell viability (data not shown). Together these data provide evidence for enhanced E-ALF-*M.tb* intracellular growth in ATs compared to bacteria exposed to A-ALF.

**Figure 1.**
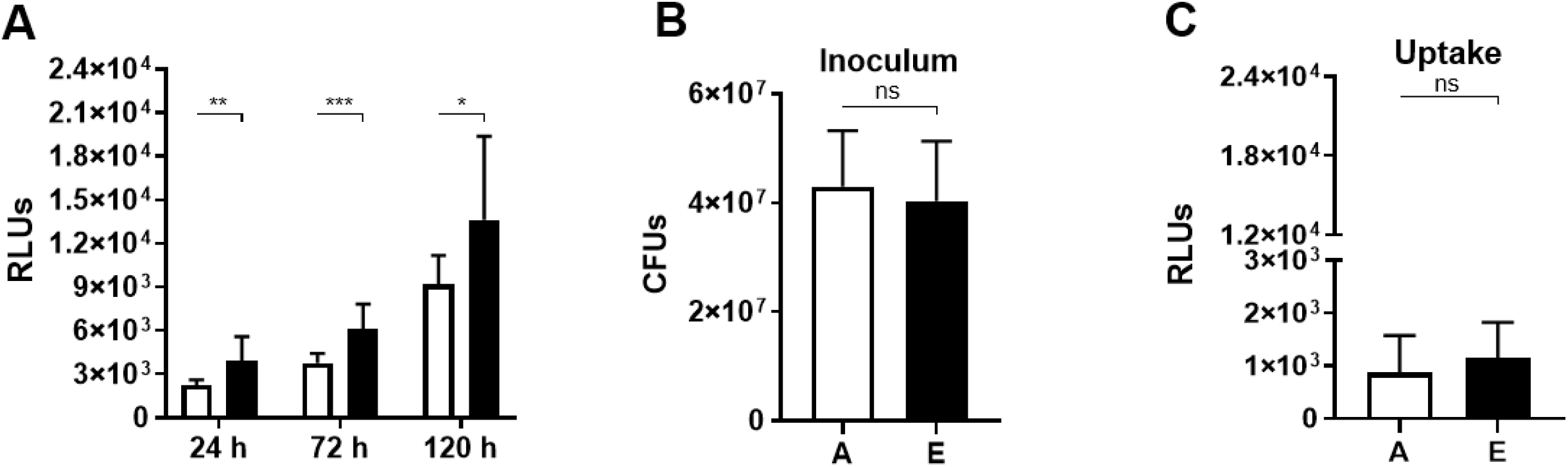
Exposure of *M.tb* to elderly human ALF is associated with increased bacterial intracellular growth in ATs. ATs were infected with ALF-exposed *M.tb* (H_37_R_v_-Lux) for 2 h at Multiplicity of Infection (MOI) of 10:1 followed by 1 h of gentamicin to kill extracellular *M.tb*. (**A**) Infected monolayers in 96-well plates were read for increased luminescence (indicative of *M.tb* H_37_R_v_-Lux intracellular growth in ATs) over time (up to 120 h), using the GLOMAX reading system. (**B**) No differences in bacteria inoculum between conditions. (**C**) No differences in bacteria uptake by ATs after 2 h post-infection. Overall data of n=4 experiment in triplicate [Mean ± SEM], using four different A-ALFs and E-ALFs. Student’s unpaired *t*-Test; Adult vs Elderly, *p<0.05, **p<0.01, ***p<0.001, ns: no significant differences. A: Adult ALF-exposed *M.tb* (white bar), E: Elderly ALF-exposed *M.tb* (black bar), RLUs: Relative Lux Units; “n” values represent the number biological replicas using a different ALF sample from different adult or elderly human donors.

We next determined the rate of bacterial replication of A-ALF *vs*. E-ALF-exposed *M.tb* during AT infection, using the fluorescent replication reporter SSB-GFP, smyc’::mCherry *M.tb* strain. This *M.tb* reporter contains a single-stranded DNA binding protein that allows for the quantification of actively replicating bacteria.^23^ Our results indicate that E-ALF-exposed *M.tb* have enhanced replication at 24 h post-AT infection (**Fig. 2**), which supports the increased growth of E-ALF *M.tb* within ATs over time (**Fig. 1A**). This increase in replication was observed until 72 h post-infection; however, at this time point, both A-ALF and E-ALF had a similar replication rate (**Fig. 2A**).

**Figure 2.**
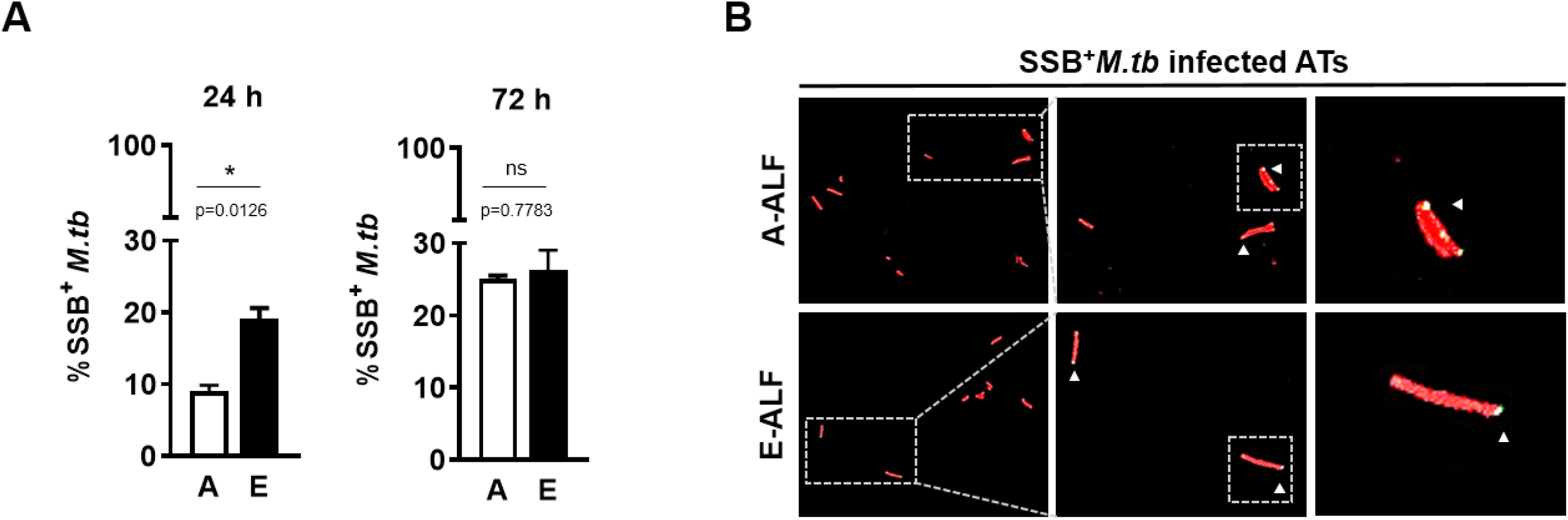
Elderly ALF-exposed *M.tb* has enhanced early replication during ATs infection. ATs were infected with the reporter SSB-GFP, smyc’::mCherry *M.tb* strain for up to 72 h, and *M.tb* replication rate was determined by confocal microscopy. (**A**) Percentage of SSB+*M.tb* exposed to A- and E-ALFs at 24 h and 72 h post-infection, n=2-4 [Mean ± SEM], using two different A-ALFs and four different E-ALFs. (**B**) Representative confocal images of ATs infected with A- and E-ALF-*M.tb* at 72 h post-infection. The region indicated by gray dashed-line is shown expanded on the right (top panels, A-ALF; bottom panels, E-ALF). Replicating SSB+*M.tb* are indicated by white arrowheads, showing merged (yellow) foci. Events were enumerated by counting at least 50 independent events, in replicates. Student’s unpaired *t*-Test; Adult vs Elderly, *p<0.05, **p<0.01, ***p<0.001, ns: no significant differences. A: Adult exposed *M.tb*, E: Elderly exposed *M.tb*; “n” values represent the number biological replicas using a different ALF sample from different adult or elderly human donors.

### E-ALF exposure decreases *M.tb* bacilli trafficking to late endosomes within Ats

Differential trafficking of *M.tb* within ATs may allow for bacterial evasion of killing or serve as a host killing mechanism,^13^ depending on cellular location. Using GFP expressing *M.tb*, we quantified co-localization of A-ALF *vs*. E-ALF-exposed *M.tb* with AT intracellular markers visualized by laser scanning confocal microscopy: Rab5 is an early endosomal marker, Rab7 is a late endosomal marker, LC3 is a marker of autophagosomes, LAMP-1 and CD63 are lysosomal markers, and ABCA1 (marker of multivesicular bodies) and ABCA3 (marker of lamellar bodies) are ATP-binding cassette lipid transporters. E-ALF-exposed *M.tb* demonstrated significantly decreased co-localization events for the late endosomal marker, Rab7, at 72 hours post-infection (**Fig. 3**). Single co-localizations of Rab5 (and not Rab7) with GFP expressing *M.tb* were not detected, consistent with previous studies in which at just 4 h post-infection nearly all *M.tb* bacilli were associated with Rab7 positive compartments.^24^ For the remainder of the intracellular markers studied (lysosome LAMP-1 and CD63, autophagosome LC3, multivesicular body ABCA1, lamellar body ABCA3), there were no significant differences in the percentage of A-ALF-exposed GFP *M.tb* co-localization events *vs*. E-ALF-exposed GFP *M.tb* co-localization events, at 72 hours post-infection (**Fig. 4**). Moreover, A-ALF or E-ALF-exposed *M.tb* were in equivalent acidified vacuoles as indicated by their co-localization with LysoTracker-Red (**Fig. 5**). Overall, exposure to E-ALF significantly decreases trafficking of *M.tb* to late endosomes within ATs.

**Figure 3.**
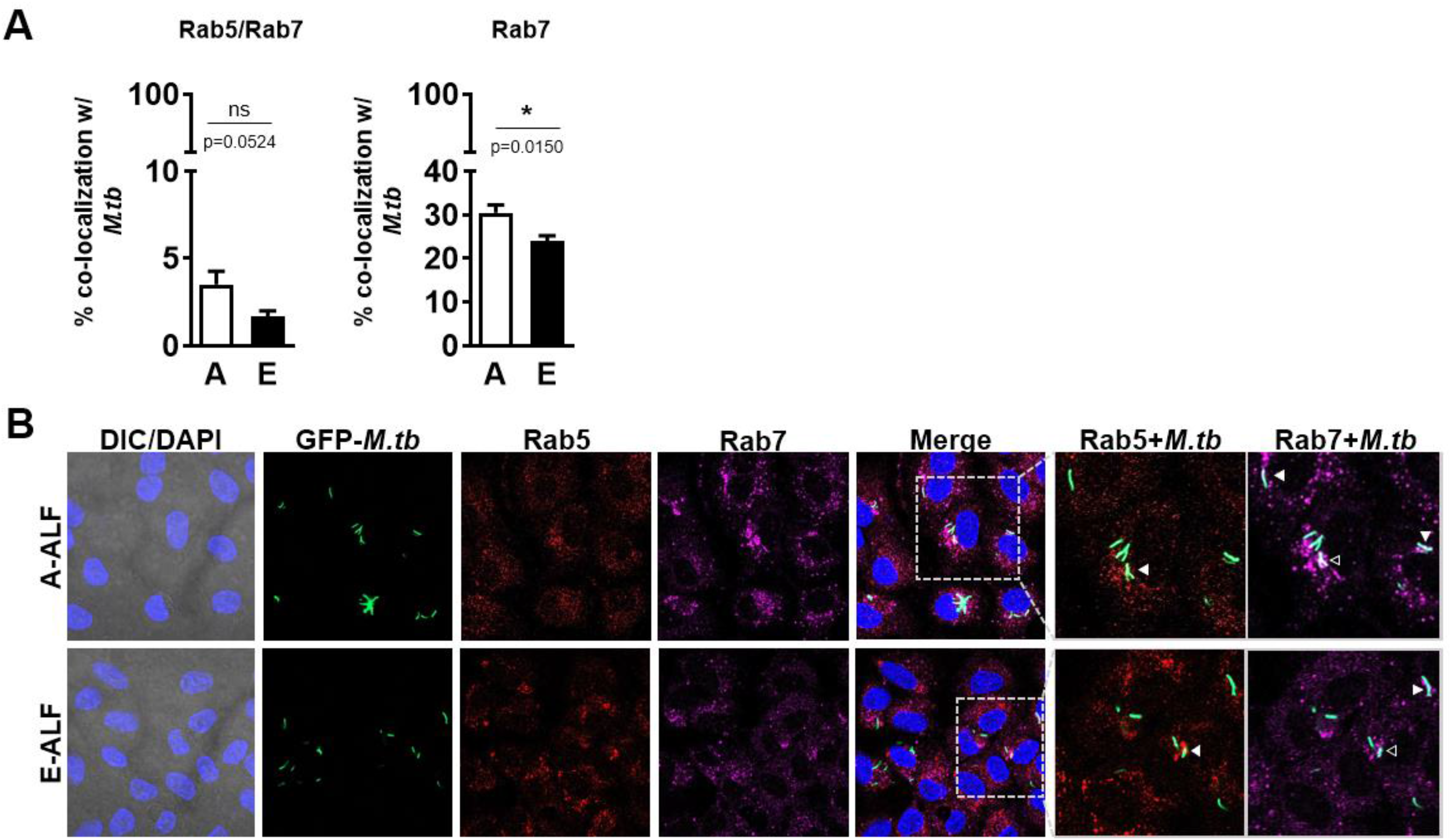
E-ALF drive decrease *M.tb* bacilli trafficking late endosomes within ATs. ATs were infected with either A-ALF or E-ALF-exposed GFP *M.tb* for 2 h at Multiplicity of Infection (MOI) of 10:1 followed by 1 h of gentamicin to kill extracellular *M.tb*. Monolayers were stained with different intracellular markers at 72 h post-infection. (**A**) Semi-quantification of Rab5+Rab7+*M.tb* (indicative of *M.tb* movement from early to late endosomes) co-localization events and Rab7+*M.tb* co-localization events (indicative of *M.tb* already in late endosomes), n=3, using three different A-ALFs and E-ALFs. (**B**) Representative confocal images of ATs infected with A- and E-ALF-*M.tb* stained with intracellular markers Rab5 and Rab7. Events were enumerated by counting at least 100 independent events, in duplicate [Mean ± SEM]. The region indicated by gray dashed-line is shown expanded on the right; and co-localization events are indicated by white arrowheads. Open arrowheads indicate double co-localization events (Rab5+Rab7+). Student’s unpaired *t*-Test; Adult vs Elderly, *p<0.05, **p<0.01, ***p<0.001, ns: no significant differences. A: Adult exposed *M.tb*, E: Elderly exposed *M.tb*, DIC: Differential Interference Contrast, DAPI: 4’,6-diamidino-2-phenylindole (ATs nuclear DNA); “n” values represent the number biological replicas using a different ALF sample from different adult or elderly human donors.

**Figure 4.**
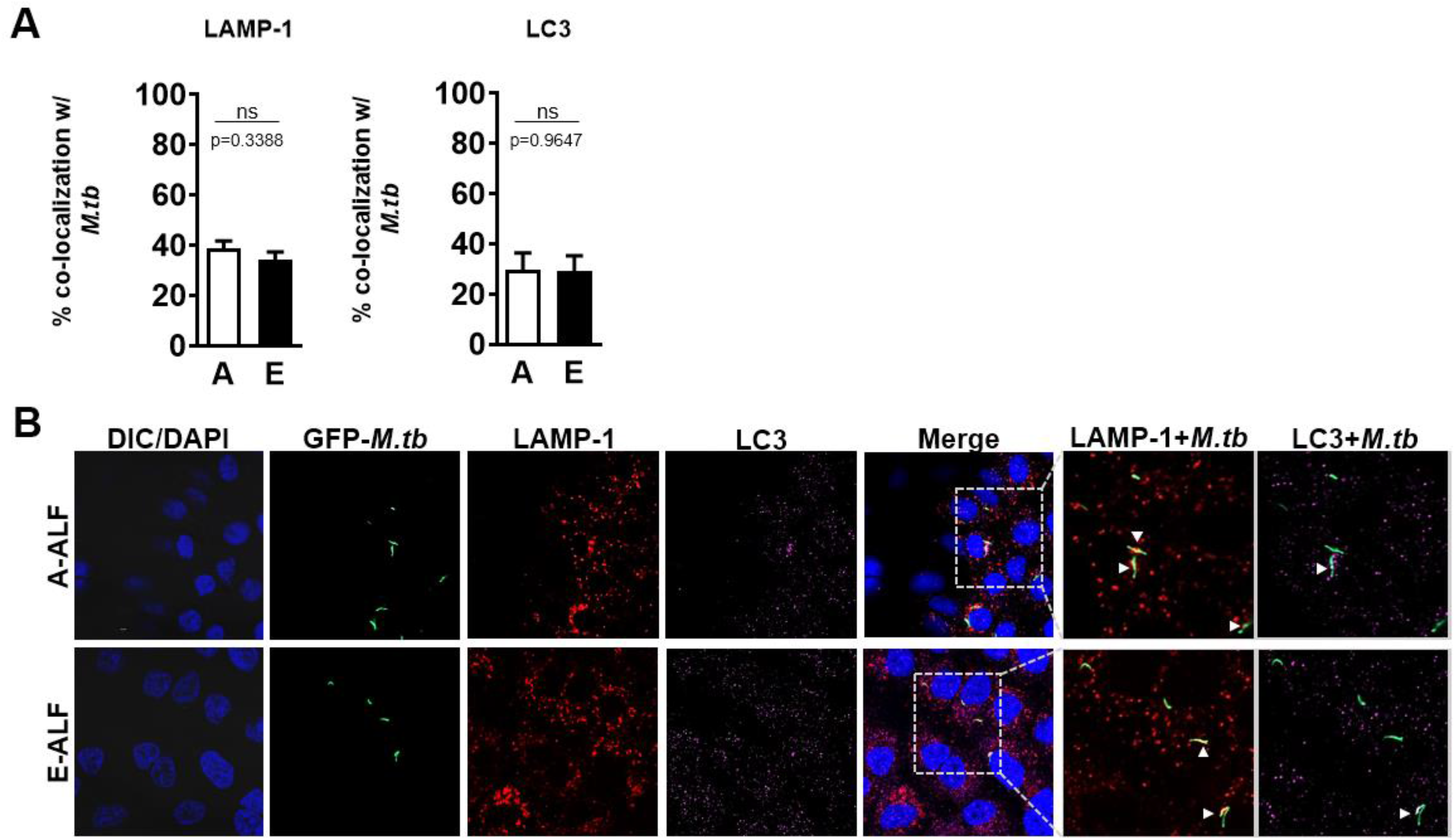
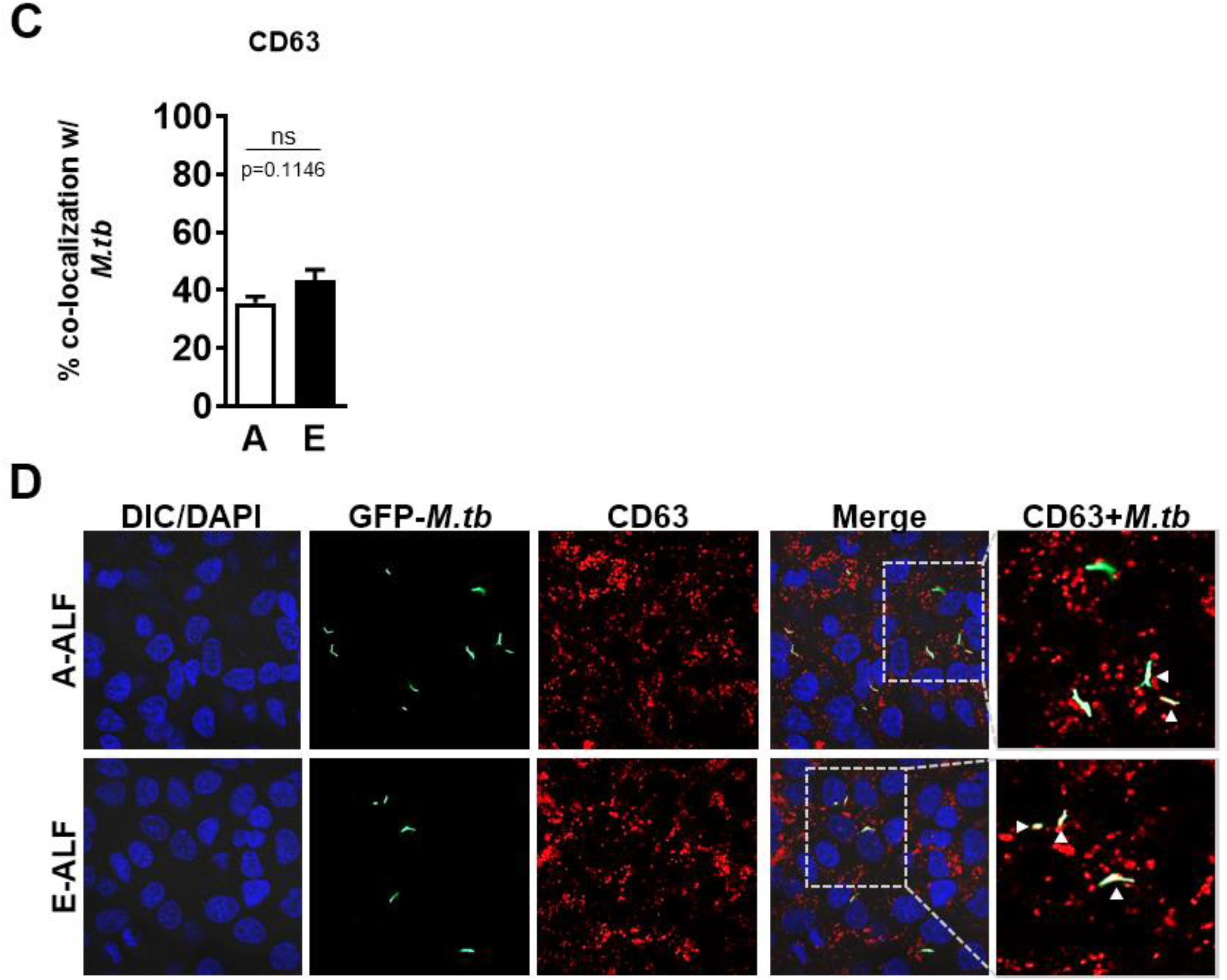
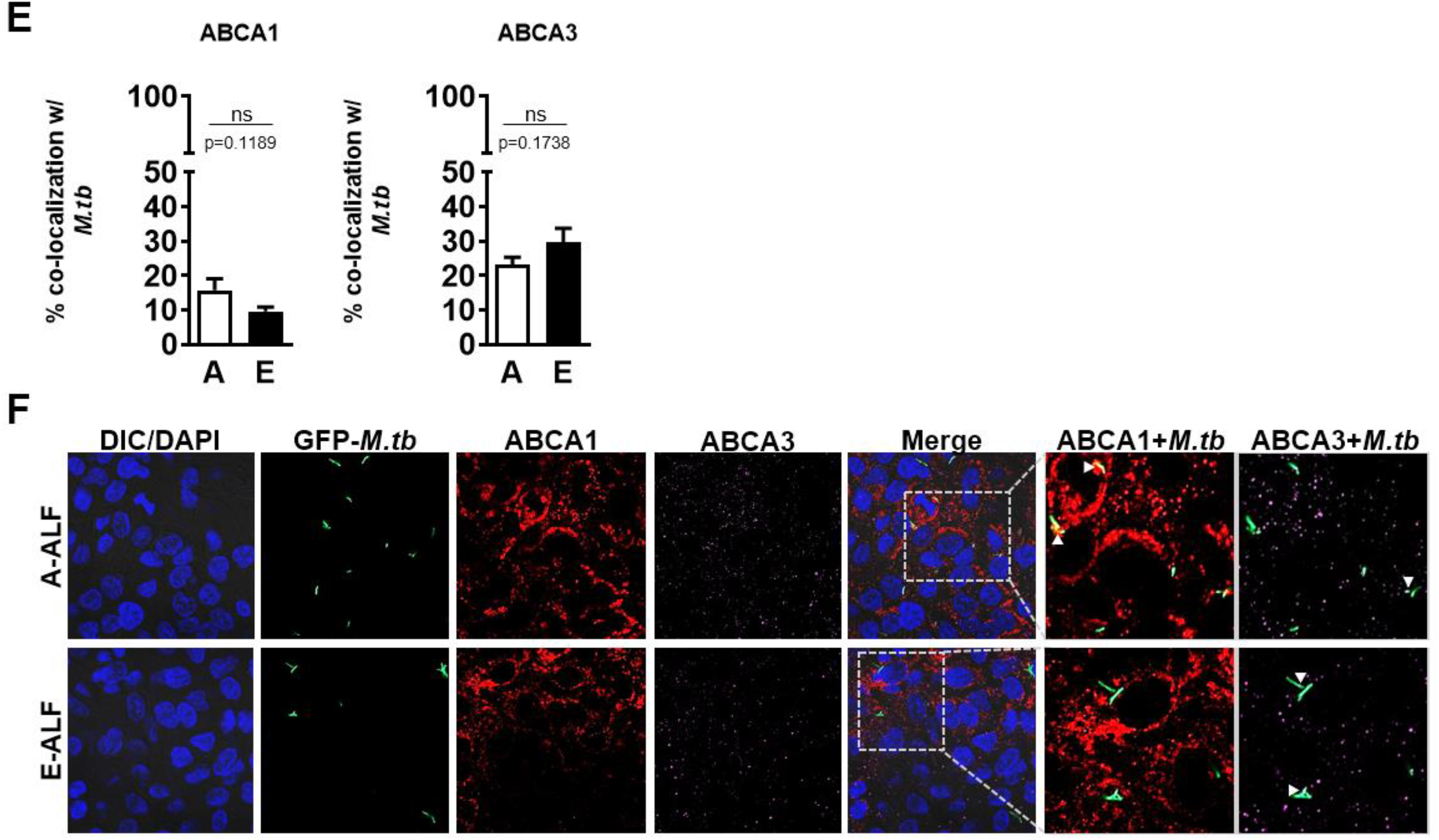
A-ALF and E-ALF does not alter *M.tb* additional intracellular trafficking within ATs. ATs were infected with either A-ALF or E-ALF-exposed GFP *M.tb* for 2 h at Multiplicity of Infection (MOI) of 10:1 followed by 1 h of gentamicin to kill extracellular *M.tb*. Stained with different intracellular markers at 72 h post-infection. (**A**) Semi-quantification of LAMP-1+*M.tb* co-localization events (indicative of *M.tb* in lysosomes) and LC3+*M.tb* co-localization events (indicative of *M.tb* in autophagosomes), n=5, using five different A-ALFs and E-ALFs. (**B**) Representative confocal images of ATs infected with A- and E-ALF-*M.tb* stained with intracellular markers LAMP-1 and LC3. (**C**) Semi-quantification of CD63+*M.tb* co-localization events (indicative of *M.tb* in lysosomes), n=3, using three different A-ALFs and E-ALFs. (**D**) Representative confocal images of ATs infected with A- and E-ALF-*M.tb* stained with intracellular marker CD63. (**E**) Semi-quantification of ABCA1+*M.tb* co-localization events (indicative of *M.tb* in multivesicular bodies) and ABCA3+*M.tb* co-localization events (indicative of *M.tb* in lamellar bodies), n=3, using three different A-ALFs and E-ALFs. (**F**) Representative confocal images of ATs infected with A- and E-ALF-*M.tb* stained with intracellular markers ABCA1 and ABCA3. Events were enumerated by counting at least 100 independent events, in duplicate [Mean ± SEM]. The region indicated by gray dashed-line is shown expanded on the right; and co-localization events are indicated by white arrowheads. Student’s unpaired *t*-Test; Adult vs Elderly, *p<0.05, **p<0.01, ***p<0.001, ns: no significant differences. A: Adult exposed *M.tb*, E: Elderly exposed *M.tb*, DIC: Differential Interference Contrast, DAPI: 4’,6-diamidino-2-phenylindole (ATs nuclear DNA); n” values represent the number biological replicas using a different ALF sample from different adult or elderly human donors.

**Figure 5.**
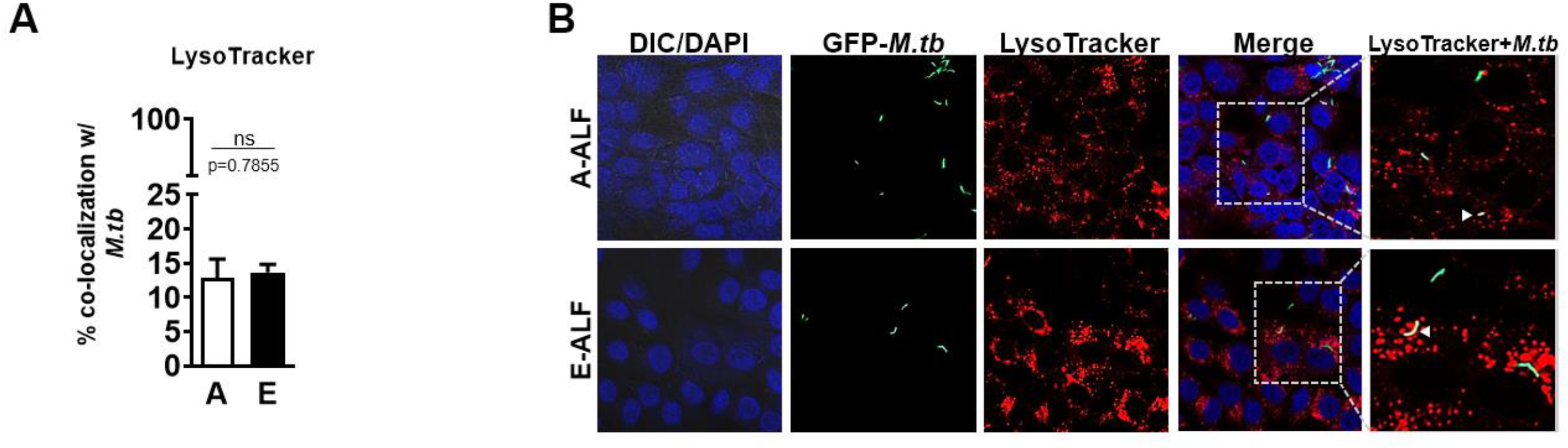
Similar acidification rates of intracellular compartments containing A- and E-ALF-exposed *M.tb*. (**A**) Semi-quantification of LysoTracker+*M.tb* acidification events, n=6 [Mean ± SEM], using six different A-ALFs and E-ALFs in infected ATs after 72 h. (**B**) Representative confocal images showing ALF-exposed GFP *M.tb* (top panels, A-ALF; bottom panels, E-ALF) co-localization with LysoTracker (red). Gray dashed-line box region is shown expanded on the right, with acidified events (yellow) indicated by white arrowheads. Events were enumerated by counting at least 100 independent events, in duplicate. Student’s unpaired *t*-Test; Adult vs Elderly, *p<0.05, **p<0.01, ***p<0.001, ns: no significant differences. A: Adult exposed *M.tb*, E: Elderly exposed *M.tb*, DIC: Differential Interference Contrast, DAPI: 4’,6-diamidino-2-phenylindole (ATs nuclear DNA); “n” values represent the number biological replicas using a different ALF sample from different adult or elderly human donors.

### E-ALF-exposed *M.tb* has increased cytosolic location in infected ATs

While it is well accepted that *M.tb* traffics through the endosomal pathway in phagocytic cells, less is known about *M.tb* intracellular localization within non-phagocytic cells such as ATs. Bacteria, including *M.tb*, can escape from phagosomes into the host cell cytosol as an alternative mechanism of survival within phagocytes.^25, 26^ Our results indicate that, at 72 hours after infection, A-ALF-exposed *M.tb* is primarily located in endosomal/lysosomal (vacuoles) compartments in ATs using transmission electron microscopy (TEM) (**Fig. 6**). In contrast, a much greater percentage of E-ALF-exposed *M.tb* was observed in the cytosol (56.3% *vs*. 16.9%) (**Fig. 6A**). The designation of cytosolic bacteria was determined by a lack of vacuolar (endosomal) membranes surrounding bacteria (**Fig. 6B**). Together, the TEM and confocal microscopy results define the overall location distribution of E-ALF and A-ALF-exposed *M.tb* within AT intracellular compartments (**Fig. 6C**), demonstrating that the majority of A-ALF-exposed *M.tb* remain in late endosomal compartments while E-ALF-exposed bacteria are increasingly present in the cytosol. We speculate that increased translocation of *M.tb* from the endosome to the cytosol for E-ALF-exposed *M.tb* may represent one mechanism by which the bacteria are able to have increased intracellular growth in ATs.

**Figure 6.**
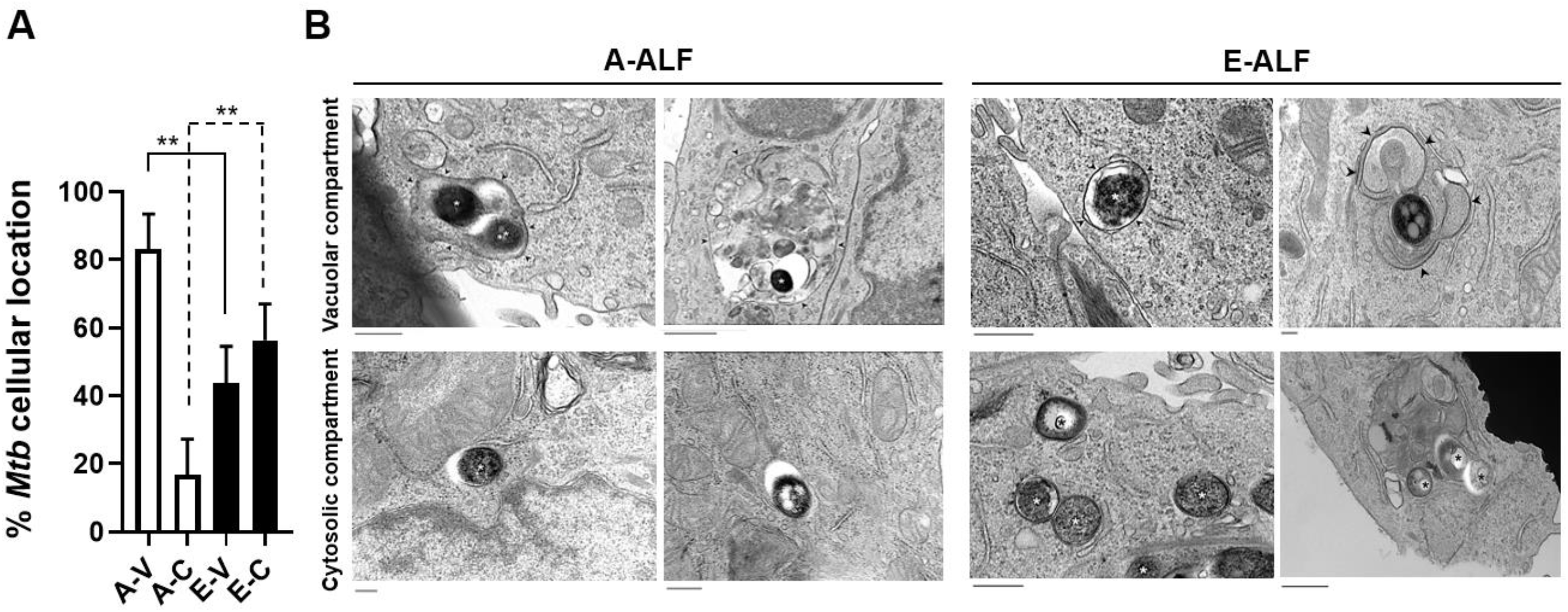
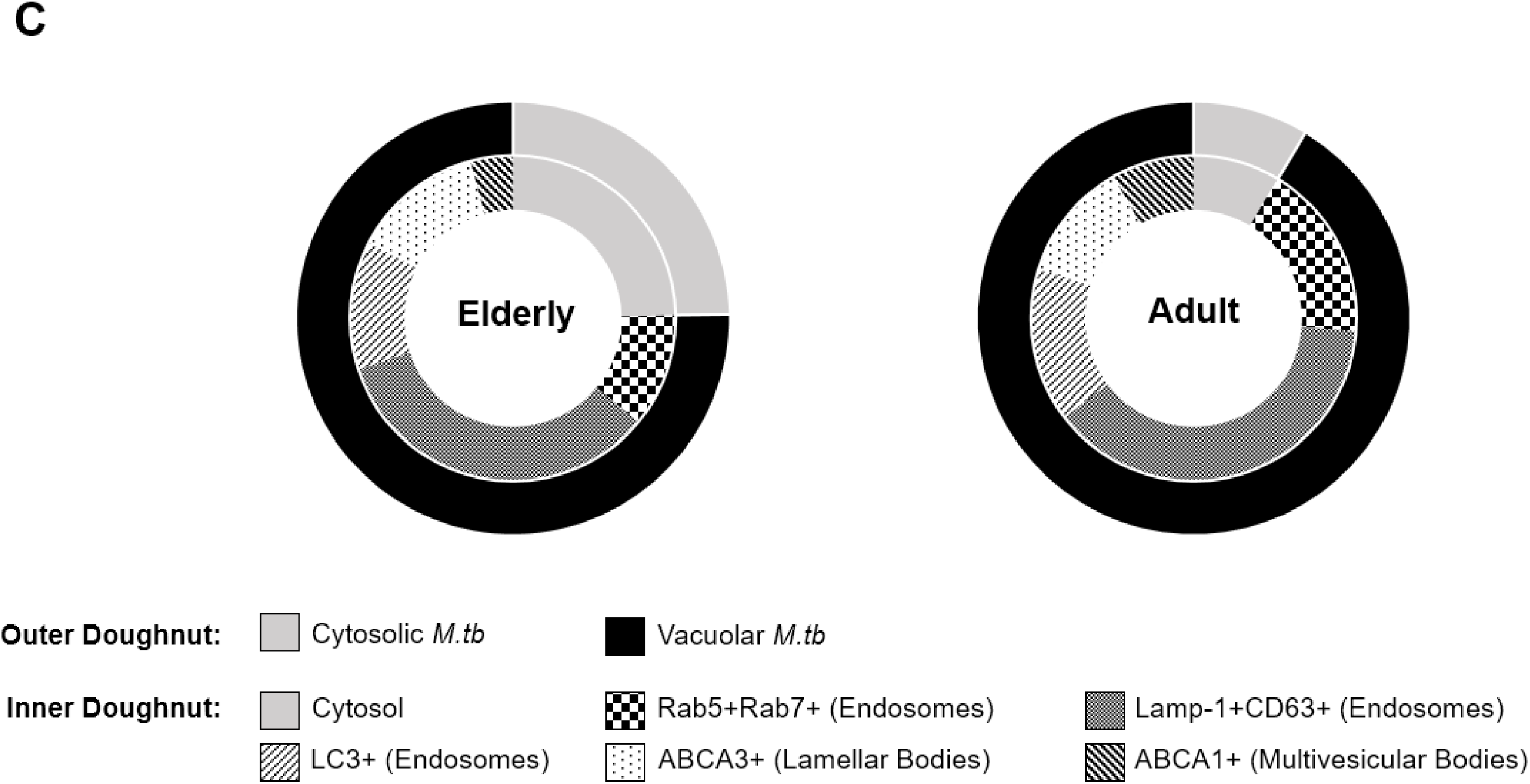
ALF-exposed *M.tb* drives differences in intracellular localization within ATs. Relative proportion of intracellular bacteria located within a membrane-bound vesicles or free in the cytosol by TEM. (**A**) Coded samples were scored by blinded analysis to quantify A-ALF- and E-ALF-exposed *M.tb* located in vacuolar (endosomal/lysosomal) or cytosolic compartments, n=2. (**B**) Transmission electron micrographs of ALF-exposed *M.tb*. Adult ALF and Elderly ALF-exposed *M.tb* were scored as “cytosolic” if they were not enclosed within a membrane, or scored as “vacuolar” if they were surrounded by a vacuolar membrane. Vacuole membranes are indicated by black arrowheads. Bacteria are indicated by asterisks. Values were determined by counting at least 100 independent events (bacteria), in replicate [Mean ± SEM]. Student’s unpaired *t*-Test; *p<0.05, **p<0.01, ***p<0.001, ns (or absence of line): no significant differences. A-V: Adult ALF-exposed *M.tb* in vacuolar compartment (Size bar 400 nm and 800 nm, respectively), A-C: Adult ALF-exposed *M.tb* in cytosolic compartment (Size bar 400 nm), E-V: Elderly ALF-exposed *M.tb* in vacuolar compartment (Size bar 600 nm and 200 nm, respectively), E-C: Elderly ALF-exposed *M.tb* in cytosolic compartment (Size bar 400 nm and 600 nm, respectively). (**C**) Normalized cellular compartment distribution of E-ALF-exposed *vs*. ALF-exposed *M.tb* within ATs; “n” values represent the number biological replicas using a different ALF sample from different adult or elderly human donors.

### Effect of A- and E-ALF-*M.tb* on AT surface marker expression

ATs are the major structural cell population of the alveolar environment^14^ and, active participants in lung immunity. ATs can activate infiltrating myeloid cells and lymphocytes by acting as antigen-presenting cells through surface expression of major histocompatibility complexes I and II (MHCI/II).^27, 28^ In this regard, at 24 h post-infection, we observed a higher percentage of HLA-ABC (MHC Class I) expression in both A-ALF and E-ALF-*M.tb*-infected ATs in comparison with HLA-DR/DP/DQ (MHC Class II) (**Fig. 7**), and this expression increased over time (120 h) (**Fig. 7A; Suppl. Fig. S2A**). Moreover, A-ALF *M.tb*-infected ATs showed an increase (compared with uninfected ATs) in the percentage of surface expression and MFI values for HLA-DR/DP/DQ over time although the data did not reach statistical significance (**Fig. 7B; Suppl. Fig. S2B**). Interestingly, HLA-DR/DP/DQ expression in E-ALF *M.tb*-infected ATs remained unchanged with respect to uninfected ATs (**Fig. 7B**), which could derive in less T cell activation in the elderly. Thus, we observed that both E-ALF and A-ALF-*M.tb* drive similar MHC surface markers expression in ATs.

**Figure 7.**
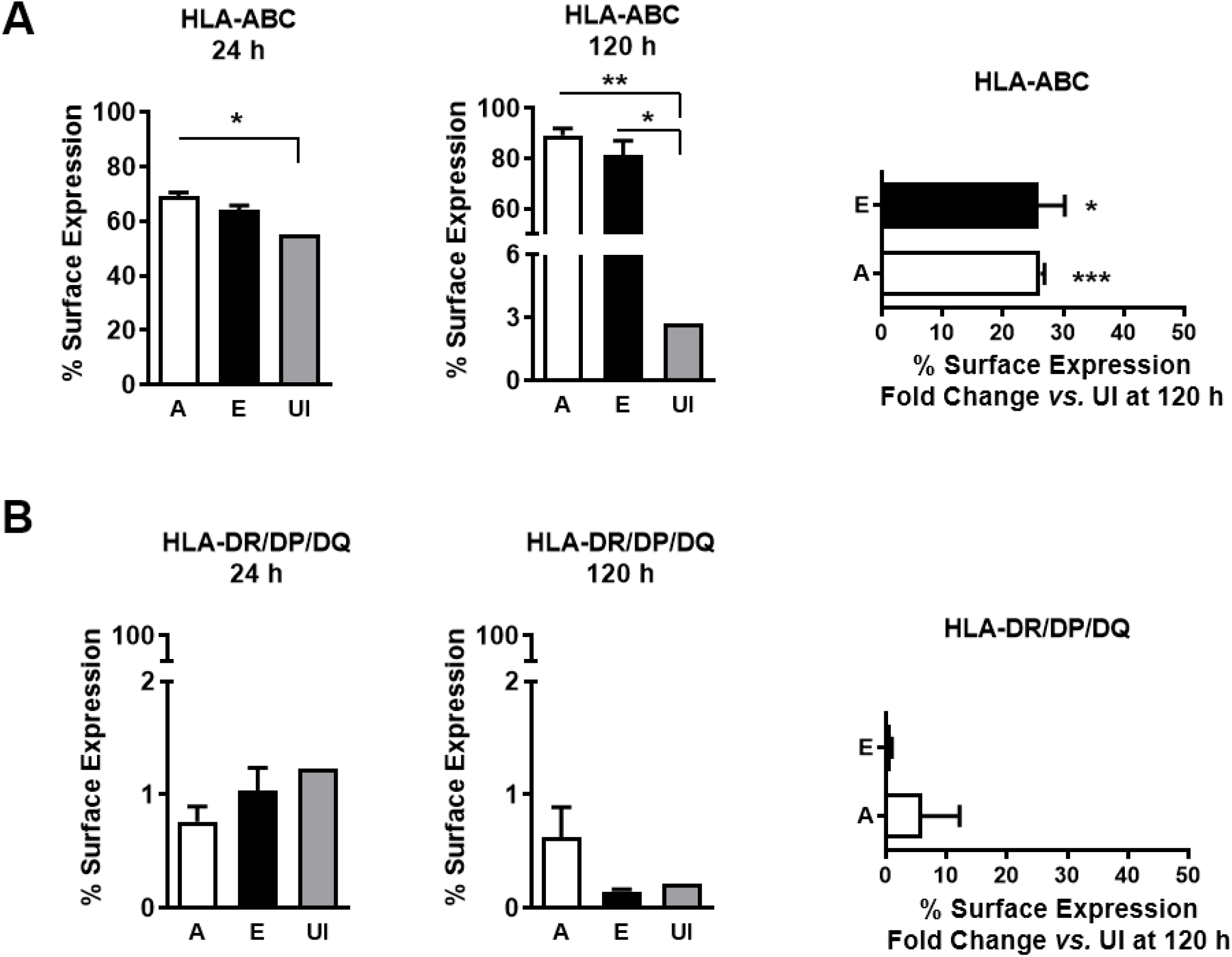
Effect of A- and E-ALF-*M.tb* on AT surface markers expression. Surface expression of uninfected and infected ATs with either A-ALF or E-ALF-exposed *M.tb* were measured by flow cytometry. (**A**) Percentage of HLA-ABC (MHC Class I) expression overtime. (**B**) Percentage of HLA-DR/DP/DQ (MHC Class II) expression overtime. Data shown are n=3 [Mean ± SEM], using three different A-ALFs and E-ALFs. Student’s unpaired *t*-Test; Adult vs Elderly, *p<0.05, **p<0.01, ***p<0.001, ns (or absence of line): no significant differences. A: Adult exposed *M.tb*, E: Elderly exposed *M.tb*, UI: Uninfected ATs; “n” values represent the number biological replicas using a different ALF sample from different adult or elderly human donors.

### Effect of A-ALF and E-ALF-*M.tb* on the production of immune mediators

Considering that *M.tb* exposure to E-ALF drives increased intracellular bacterial growth in ATs, we tested whether E-ALF-exposed *M.tb* increase levels of AT pro-inflammatory cytokines and chemokines, reflecting increased AT activation. We quantified the production of immune mediators responsible for immune cell infiltration toward the site of infection and/or for promoting immune cell proliferation and maturation in the AT cultures.^29^ Both A-ALF and E-ALF-exposed *M.tb* infection induced mainly pro-inflammatory cytokines by ATs when compared to uninfected ATs, which was maintained over time (up to 120 h studied). This increasing trend was the case for TNF and IL-6, with IL-18 reaching significant levels (**Fig. 8A**). Significant production of chemokines was also observed from AT cultures infected with A-ALF and E-ALF-exposed *M.tb* compared to uninfected ATs. This was particularly the case for the following chemokines: CCL2/MCP-1, CCL5/RANTES, and IL-8/CXCL8 at later time points post-infection (**Fig. 8B**). GM-CSF also was significantly produced at later time-points (**Fig. 8B**). Overall, we did not observe significant differences in AT immune mediator production during infection between A-ALF or E-ALF-exposed *M.tb*. Thus, exposure to E-ALF enhances *M.tb* replication and growth in ATs without altering cell activation compared to A-ALF-exposed bacteria.

**Figure 8.**
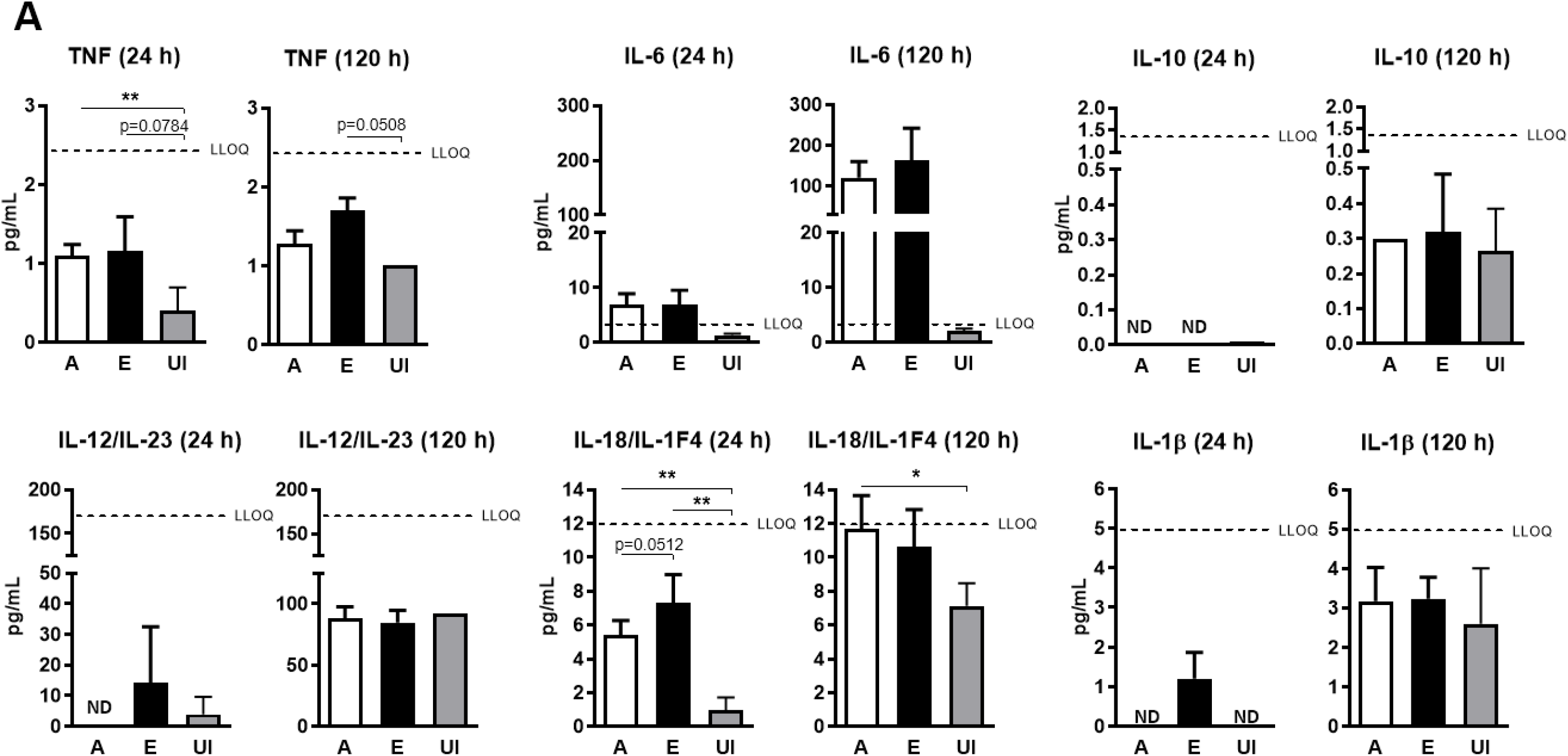
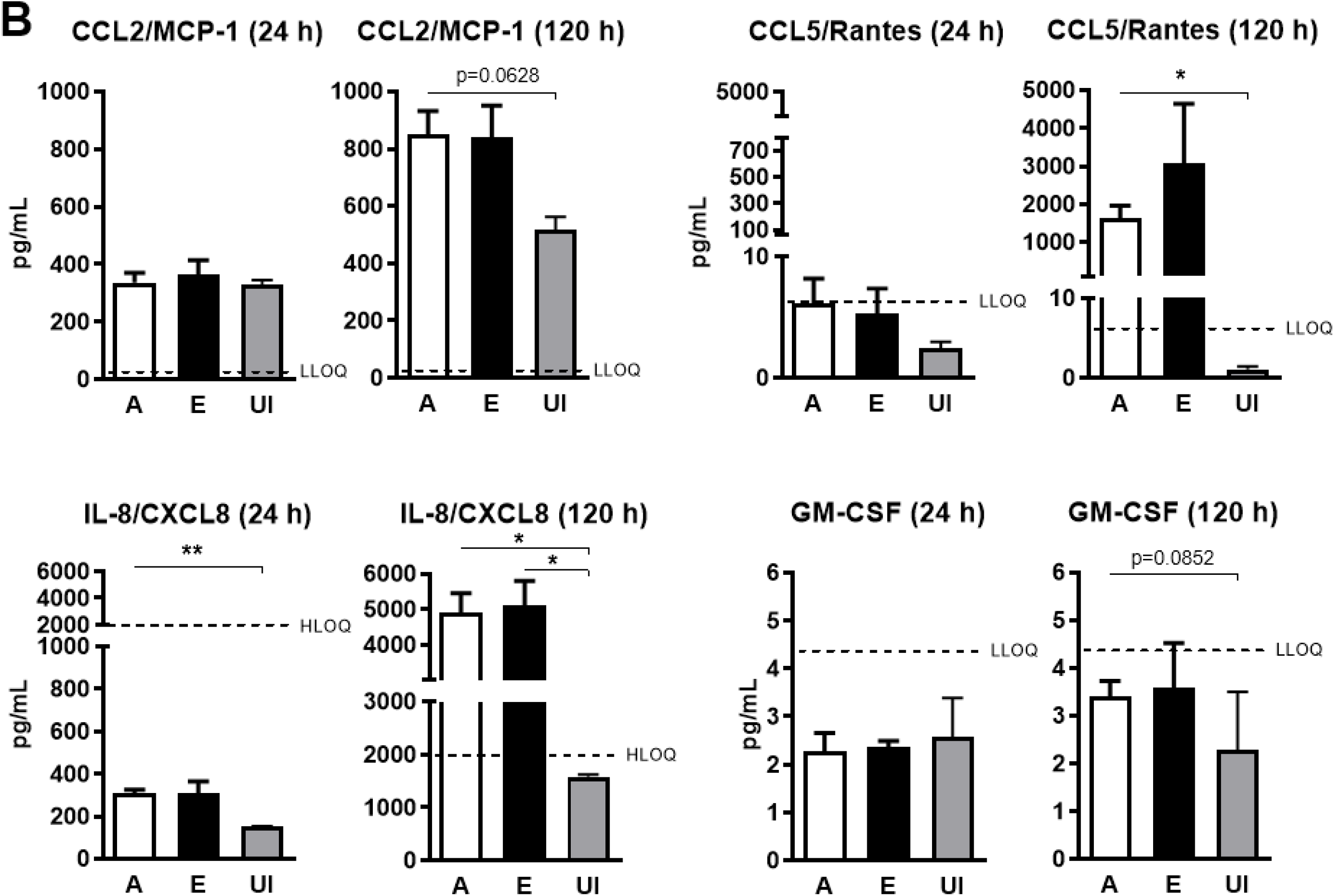
Effect of A- and E-ALF-*M.tb* on the production of ATs immune mediators. ATs supernatants from uninfected and infected ATs with either A-ALF or E-ALF-exposed *M.tb* were tested for cytokines (**A**) and chemokines (**B**) production and measured over time by Luminex multiplex assay following the manufacturer’s instructions. Cell supernatants tested were undiluted and the data shown are n=5 (using five different A-ALFs and E-ALFs) and n=2 (uninfected ATs conditions). For CCL-2/MCP-1 and IL-8/CXCL8 cell supernatants were diluted 1:15 and 1:5, respectively, and measured over time by ELISA kits following the manufacturer’s instructions. Student’s unpaired *t*-Test [Mean ± SEM]; Adult vs Elderly, *p<0.05, **p<0.01, ***p<0.001, ns (or absence of line): no significant differences. A: Adult exposed *M.tb*, E: Elderly exposed *M.tb*, UI: Uninfected ATs, ND: Not detectable, LLOQ: Low limit of quantification, HLOQ: High limit of quantification.

## DISCUSSION

TB remains one of the top 10 causes of death worldwide.^1^ The elderly population (65 years or older) with inherent compromise in immunity is at higher risk of developing active TB disease.^3^ Additionally, the elderly may have additional comorbidities (*e.g*., diabetes, hyperglycemia, HIV co-infection, malnutrition, smoking, among others) that would make them even more vulnerable to primary *M.tb* infection and reactivation of a latent infection to active TB disease.^3, 30, 31^ Therefore, it is critical to study how the lung environment changes as we age, and the impact of these changes on the establishment of respiratory infections, such as *M.tb*. We have published that the elderly lung mucosa contains many oxidized proteins and constitutes an inflammatory environment, with dysfunctional surfactant proteins and complement function.^9^ Moreover, we have established that upon contacting the elderly lung mucosa, *M.tb* replicates faster in human macrophages and has increased bacterial burden *in vivo* in C57BL/6 mice, inducing increased lung tissue damage.^9^ Here we determined that after exposure to the elderly lung mucosa or E-ALF, *M.tb* infects alveolar epithelial cells (ATs), the major non-phagocytic cell of the alveolar space, equivalently to A-ALF-exposed *M.tb*. However, E-ALF exposed bacteria replicated faster inside ATs. This finding was associated with the fact that E-ALF-exposed bacteria were found to a much greater extent in the cytosol, suggesting that this location is a favorable niche for the bacteria to establish infection, averting host immune response (**Fig. 9**).

**Figure 9.**
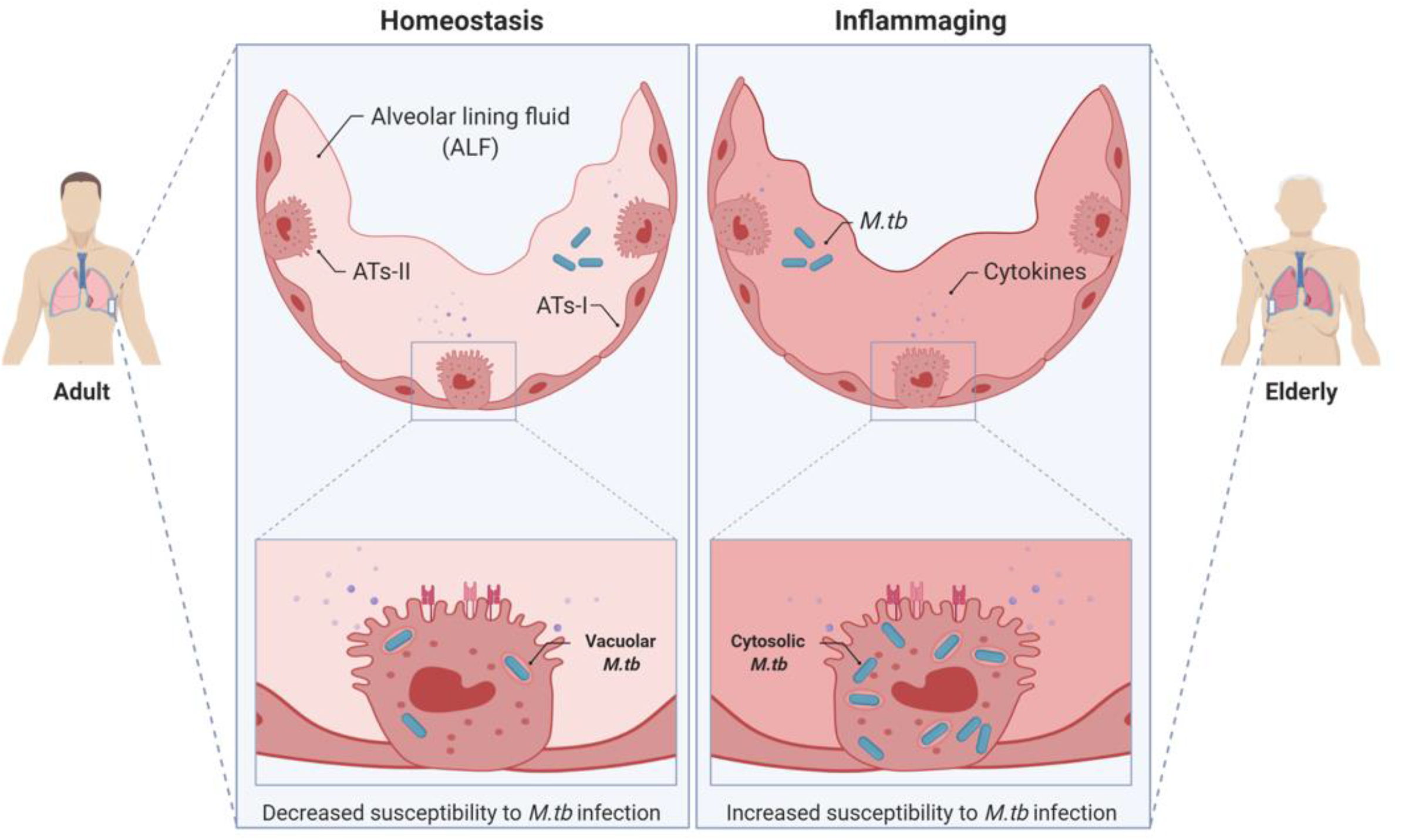
Schematic overview of the main findings in this study. *Mycobacterium tuberculosis* (*M.tb*) exposure to alveolar lining fluid (ALF) from elderly individuals enhances intracellular growth in alveolar epithelial cells (ATs). Moreover, ATs infection with E-ALF-exposed *M.tb* does not show altered production of inflammatory mediators (cytokines and chemokines) or cell activation. A-ALF-exposed *M.tb* is mainly located in endosomal/lysosomal (vacuoles) controlling its growth, but interestingly, E-ALF-exposed *M.tb* appears to be located in both vacuolar and cytosolic compartments. Overall, E-ALF seemingly promotes *M.tb* growth within ATs, preventing cell activation, and potentially exploiting the AT cytosol as a niche for survival. This illustration was created with BioRender (https://biorender.com/).

Once *M.tb* reaches the alveolar space, it is recognized by resident or recruited phagocytes, including alveolar macrophages, dendritic cells, and/or neutrophils.^32^ Although *M.tb* uptake by phagocytes does not always lead to clearance of infection, macrophages from elderly individuals are more permissive to intracellular growth.^2, 33^ Another cell population particularly crucial in the early stages and outcomes of *M.tb* infection are the non-professional phagocytes ATs.^15, 34^ Ats line the alveolar epithelium forming a physical barrier that prevents rapid invasion and plays a critical role in host defense to control *M.tb* infection.^15, 32^ One of the significant roles is the production of components within the alveolar mucosa, or alveolar lining fluid (ALF), composed of a surfactant monolayer and an aqueous hypophase surrounding alveolar compartment cells.^15^ We have shown that innate soluble components of ALF, including homeostatic hydrolytic enzymes, modify the *M.tb* cell envelope and allow for better recognition by cells of the immune system.^4-6^ In this regard, it remains unclear if E-ALF-exposed *M.tb* infection of ATs alters their production of ALF innate soluble components, which would also impact the outcome of infection in the elderly.

Our group has previously published that ALF from elderly individuals have increased levels of pro-inflammatory and pro-oxidative mediators.^9^ Furthermore, human macrophages infected with elderly human ALF–exposed *M.tb* had reduced control of infection.^9^ Considering that most studies focus only on the role of ALF in altering *M.tb*-phagocyte interactions, we aimed to determine the impact of the adult and elderly lung mucosa on *M.tb* infection of ATs. Here, we show that *M.tb* exposure to human E-ALF drove significantly increased intracellular bacterial growth in ATs. This finding supports the importance of ALF in old age which has a different composition^7^ in enhancing *M.tb* infection by allowing *M.tb* to replicate faster in both alveolar phagocytic^9^ and non-phagocytic cells. Our results also indicate that E-ALF-exposed *M.tb* does not activate ATs; nor kill them, enabling the continued growth of E-ALF-exposed *M.tb* within ATs over time.

The *M.tb* phagosome in macrophages shares features with early endosomes due to blockage of EEA1 and Rab7 recruitment,^35, 36^ whereas *M.tb* was found to traffic to late endosomes in epithelial cells.^37^ Our results are consistent with previous studies in which *M.tb* vacuoles in alveolar and bronchial epithelial cell lines resides mainly in Rab7 positive compartments.^24, 37^ Notably, exposure to E-ALF decreased *M.tb* endosome association with Rab7 in late endosomes in ATs. In this regard, we found similar results in ATs infected with higher MOIs (data not shown). Our data showed no significant differences in the percentage of co-localization events of A-ALF or E-ALF-exposed *M.tb* within additional AT intracellular compartments (*i.e*., LAMP-1, CD63, LC3, ABCA1, ABCA3) consistent with our previous studies showing that *M.tb* bacilli exposed to different human A-ALFs did not have altered intracellular trafficking in those compartments within ATs.^13^ Here we further explored the location of *M.tb* within ATs by TEM to better understand the differences in intracellular growth and trafficking. We found that E-ALF-exposed *M.tb* are located in both vacuolar and cytosolic compartments, whereas A-ALF-exposed *M.tb* bacilli are located mainly in vacuoles, where ATs could potentially better control their replication and growth.

Several studies performed in macrophages have shown how *M.tb* can employ different mechanisms of survival, including, inhibition of phagosome-lysosome fusion,^38^ suppression of the autophagy pathway^39^ and escape from phagosomes into the cytosol.^25, 26^ However, trafficking patterns of *M.tb* in ATs seem to differ as *M.tb* is contained mainly in late (Rab7) endosomal compartments and lysosomal fusion with late endosomes is inhibited, supporting its increased survival within ATs.^37^ In this regard, we note that *M.tb* bacilli traffic to late endosomes in ATs, but upon exposure to E-ALF, *M.tb* decreased association with Rab7 positive compartments as it translocated into the cytosol, a potential niche to increase its replication rate.

The autophagy pathway is an alternative killing mechanism against intracellular pathogens when they escape from the typical phagosome/lysosome fusion killing mechanism.^40, 41^ Since E-ALF-exposed *M.tb* was located in the cytosol where there is a high bacterial replication rate, we explored if autophagy was attenuated in ATs infected with E-ALF-exposed *M.tb*. Our results indicate that A-ALF and E-ALF exposure did not alter *M.tb* bacilli trafficking to autophagosomes (no differences in LC3 positive compartments) within ATs. Although autophagy has been described as an alternative killing mechanism of *M.tb* in macrophages, it could also play an opposite role in mycobacterial trafficking in ATs if it fails to eliminate the bacterial burden.^37^ In fact, inactivation of the autophagy pathway using 3-methyladenine decreased *M.tb* intracellular growth and was advantageous for ATs survival.^37^ Lastly, exclusively pathogenic mycobacteria species, including *M.tb*, are reported to translocate from the phagolysosome into the cytosol facilitated by the ESAT-6 secretion complex-1 (ESX-1 type VII secretion system) in macrophages.^42^ While more studies are needed to elucidate whether the *M.tb* ESX-1 type VII secretion system also mediates *M.tb* translocation into the cytosol in non-professional phagocytes, *M.tb* genes encoding ESAT-6 proteins are upregulated during *M.tb* infection of ATs.^43^ Given that E-ALF exposure promoted *M.tb* growth within ATs and greater translocation to the cytosol we speculate this may be a mechanism for better survival of *M.tb* in ATs. Ongoing studies in our lab are focused on determining the *M.tb* metabolic status within ATs at the time of its translocation into the cytosol to elucidate the bacterial mechanism involved in this process.

Additional mechanisms that ATs may use to contribute to the innate and adaptive immune responses after *M.tb* infection are, first, bacterial uptake by recognition and binding of microbial associated molecular patterns present on the *M.tb* surface;^32^ and second, modulation of the cell to cell crosstalk by the production of immune mediators.^29, 34^ We have found that *M.tb* infected ATs secrete some cytokines (*e.g*. IL-18), but mainly chemoattractants (*e.g*. CCL2, CCL5/RANTES, and IL-8) at later time points post-infection. Some of these immune mediators are significantly induced by E-ALF-exposed *vs*. A-ALF-exposed *M.tb* (*e.g*. CCL5/RANTES) and are critical to the initiation of local inflammatory responses, mainly cell recruitment (e.g. neutrophils), and activation of host innate cells and even induction of cell-mediated immunity after *M.tb* infection, which could unbalance local immunity towards favoring *M.tb* infection spread being detrimental for the host.^34^

Here we show that during infection in ATs, E-ALF-exposed *M.tb* induced similar changes in surface marker expression or in the production of immune mediators when compared to A-ALF-exposed *M.tb*, possibly hindering alveolar compartment cells activation, infiltration, and differentiation. Indeed, when compared to uninfected cells, E-ALF-exposed *M.tb* did not induce the expression of MHC-II on the AT surface, potentially negatively influencing T cell activation. Altogether, our data reveal that *M.tb* exposure to the inflammatory E-ALF environment which contains an array of oxidized proteins^7, 9^ enhances *M.tb* intracellular growth in ATs, while promoting translocation of bacteria to the AT cytosol as a potential niche for establishing and propagating *M.tb* infection. Our study highlights the impact of elderly lung mucosa on *M.tb* infection of ATs, critical non-professional phagocytes that impact TB as well as other respiratory infectious diseases.

## MATERIALS AND METHODS

### Ethics Statement and Human Subjects

Human subject studies were carried out in strict accordance with the US Code of Federal and Local Regulations (OSU Institutional review board (IRB) numbers 2012H0135, and 2008H0119 and Texas Biomedical Research Institute/UT-Health San Antonio/ South Texas Veterans Health Care System IRB number HSC20170673H. Bronchoalveolar lavage fluid (BALF) from healthy adults (aged 18-45 years) and elderly (aged 60 years and older) individuals were recruited from both sexes without discrimination of race or ethnicity after informed written consent. See Supplementary Information for details.

### Human ALF isolation

ALF was obtained and concentrated from human bronchoalveolar lavage fluid (BALF) from healthy adults and elderly donors. ALF was normalized as previously described,^4, 9, 11^ to obtain the physiological concentration present within the lung (at 1 mg/mL of phospholipid). Briefly, BAL was performed in sterile 0.9% NaCl, filtered (0.2 μm pore size sterile filter system), and subsequently concentrated 20-fold by using a 10-kDa molecular mass cutoff membrane Centricon Plus (Amicon Bioseparations) device at 4°C, aliquoted in low protein binding-sterile tubes and stored at −80°C.

### AT culture

For all experimental procedures, we utilized the human ATs type II-like cell line A549 (ATCC® CCL-185™) as a lung carcinoma cell line that exhibits most AT type II cells traits (model of ATs). Cell cultures were prepared as we previously described^13^ with minor changes. Briefly, the A549 cell line was cultured at 37 °C with 5% CO_2_ in DMEM/F12 supplemented with 10% FBS (Atlas Biologicals, Fort Collins, CO) and 1% PenStrep (Sigma, St. Louis, MO), allowing at least three weeks of passages prior to use in experiments. Cells were maintained in antibiotic-free growth medium one week before doing *M.tb* infection.

### M.tb cultures

*M.tb* H_37_R_v_-Lux (27294) (kindly provided by Drs. Abul Azad and Larry Schlesinger, Texas Biomedical Research Institute) was grown as described.^44, 45^ SSB-GFP, smyc’::mCherry *M.tb* Erdman (kindly provided by Dr. David Russell, Cornell University) was grown as described.^23^ GFP-*M.tb* Erdman (kindly provided by Dr. Marcus Horwitz, UCLA) was grown as previously described.^4^ Single bacterial suspensions were prepared as we described.^9, 13^

### Exposure of M.tb bacteria to human ALF

Preparation of adult and elderly ALF-exposed *M.tb* was performed as we previously described.^10, 11, 13^ ALF-exposed *M.tb* inoculums were serially diluted in 7H9 broth and plated on 7H11 agar to determine *M.tb* viability and specifically to confirming no differences in viable bacterial counts among E-ALF *vs*. A-ALF-exposed *M.tb*.

### M.tb infection and luciferase-based intracellular growth of Ats

*M.tb* infection of ATs was performed as previously described.^13^ Briefly, single cell suspensions of *M.tb* in DMEM/F12/FBS media were added to the ATs culture at various MOIs, and cells were incubated for 2 h with the first 30 min on a platform shaker. After infection, unbound bacteria were removed by washing, and gentamicin (50 μg/mL)-supplemented medium was added for 1 h to kill extracellular bacteria. Following this, cells were washed and incubated with 10 μg/mL gentamicin-supplemented medium for the indicated times. For luciferase-based *M.tb* growth assays,^45^ ATs were infected with *M.tb* H_37_R_v_-Lux, and bacterial bioluminescence was measured every 24 h for up to 120 h with a GloMax Multi Detection System (Promega, Madison, WI). For the *M.tb* replication rate experiment, ATs were infected with SSB-GFP, smyc’::mCherry *M.tb* Erdman strain. For intracellular trafficking, acidification assay, and electron microscopy experiments, ATs were infected with GFP-*M.tb* Erdman (GFP, 488 nm) strain.

### AT cell viability assay

At indicated times post-infection, AT cytotoxicity was determined by CellTiter-Glo® luminescent cell viability assay (Promega Cat. #G7570) following the manufacturer’s instructions. See Supplementary Information for details.

### Immunocytochemistry and Confocal Microscopy

ATs monolayers on glass coverslips were infected for 2 h with A-ALF– or E-ALF-exposed *M.tb* at MOI 10:1 and processed as described.^13^ Briefly, at 72 hours post-infection, ATs were fixed with cold 4% paraformaldehyde for 15 min at room temperature and permeabilized with 0.1% Triton-X100 in PBS for 10 min at room temperature. To evaluate the *M.tb* intracellular trafficking, cellular compartments were stained with primary Abs in confocal blocking buffer (5 mg/ml BSA, 5% HI-FBS, 10% donkey serum, 0.03% Triton X-100) for overnight incubation at 4°C. Following this, the secondary Abs or matched isotype controls were incubated for 1 h at 37°C.

Intracellular markers used were Rab5A (early endosomal marker; mouse anti-human Rab5A; 1:200; Cell Signaling Technology Cat. #46449), Rab7 (late endosomal marker; rabbit anti-human Rab7; 1:100; Cell Signaling Technology Cat. #9367), LAMP-1 (lysosomal marker; mouse anti-human LAMP-1; 2 μg/ml; DSHB Cat. #P11279), CD63 (lysosomal marker; mouse anti-human CD63; 1 μg/ml; BD Biosciences Cat. #556019), LC3-II (autophagy marker; rabbit anti-human LC3; 1:200; Cell Signaling Technology Cat. #3868), ABCA1 (ATP-binding cassette lipid transporter marker of multivesicular bodies; mouse anti-human ABCA1; 5 μg/ml; Abcam Cat. #ab18180) and, ABCA3 (lamellar bodies ABC transporter marker; rabbit anti-human ABCA3; 2 μg/ml; Abcam Cat. #ab99856). Secondary Abs were donkey anti-rabbit IgG H&L conjugated to Alexa Fluor® 647 (Abcam Cat. #ab150063) and donkey anti-mouse IgG H&L conjugated to Alexa Fluor® 568 (Abcam Cat. #ab175700). Negative controls were included to check for non-specific binding and false-positive results, including wells incubated with isotypes control antibodies or in which primary Abs were omitted. Isotype controls were mouse IgG1 isotype (Abcam Cat. #ab91353) and rabbit IgG isotype (Invitrogen Cat. #26102). The nucleus was stained with 50 ng/ml 4’,6-diamidino-2-phenylindole (DAPI) (Invitrogen Cat. #D1306) for 10 min at room temperature. After multiple washes to remove the excess of DAPI solution, coverslips were mounted on slides using ProLong Gold Antifade Reagent (Invitrogen Cat. #P36934).

Cells were visualized by laser scanning confocal microscopy using ZEISS LSM 800 microscope set at appropriate parameters and a final magnification of 600X. Co-localization events of the different cellular compartments containing GFP-*M.tb* Erdman was quantified by counting at least 100 events per condition in duplicate. All microscopy data were analyzed with Zeiss ZEN Software.

### AT compartment acidification assay

AT compartment acidification was determined by LysoTracker-Red® assay (Invitrogen Cat. #L7528), at 72 hours post-infection following the manufacturer’s instructions. See Supplementary Information for details.

### Transmission electron microscopy

ATs monolayers were infected for 2 h with A-ALF-exposed *M.tb* or E-ALF-exposed *M.tb* at MOI 100:1 and processed as mentioned previously. Infected ATs, at 72 hours post-infection were fixed in 2.5% glutaraldehyde and 2% formaldehyde (in 0.1 M Na Cacodylate pH 7.3) and analyzed by transmission electron microscopy (TEM) as previously described^46, 47^ with minor changes. See Supplementary Information for details.

Samples were shipped to Dr. Daniel L. Clemens, an expert in the study of intracellular compartmentalization of *M.tb* by TEM for analyses in a blinded manner.^48-50^ The main objective was to quantify the relative proportion of bacilli located within membrane-bound vesicles or located in the cytosol compartment (lacking membrane bilayers around the bacilli). For each condition, at least 100 events were imaged and scored in a blinded fashion using the JEOL 100CX transmission electron microscope (BRI Electron Microscopy Core Facility; Brain Research Institute UCLA).

### Expression of AT surface markers by flow cytometry

Following infection at different time points, ATs were stained with LIVE/DEAD (L/D) Fixable Dead Cell Kit (Invitrogen Cat. # L23105) following manufacturer’s instructions, and a panel of surface markers or matched isotype controls in deficient RPMI buffer, for 10 min in the dark at 4°C. Recombinant fluorescently-labelled Abs used were VioBlue HLA-ABC (1:50; REAfinity™ Cat. #130-120-435), PE HLA-DR/DP/DQ (1:50; REAfinity™ Cat. #130-120-715), APC CD324 [e-Cadherin] (1:50; REAfinity™ Cat. #130-111-840); and isotypes controls were VioBlue human IgG1 (1:50; REAfinity™ Cat. #130-113-442), PE human IgG1 (1:50; REAfinity™ Cat. #130-113-438) and, APC human IgG1 (1:50; REAfinity™ Cat. #130-113-434). Cells were then washed, fixed with 4% PFA, and follow by consequent washes before stored overnight at 4°C until analysis. Samples were analyzed using a FACSymphony A3 flow cytometer (BD Biosciences). Each sample for analysis contained 5,000–100,000 events, and dead cells were subsequently gated out according to their L/D staining. All flow cytometry data were analyzed using FlowJo Version 10.6.2 software (FlowJo, LLC).

### AT cytokine and chemokine production

Protein levels in supernatants from infected (A-*M.tb* and E-*M.tb*) or non-infected ATs were determined using a multiplex panel human magnetic bead Luminex® Assay for IL-6, CCL5/Rantes, GM-CSF, TNF-α, IL-1β, IL-18/IL-1F4, IL-10 and IL-12/IL-23 (R&D, Human 10-Plex Cat. #LXSAHM-10, Lot #L134898); and using Human enzyme-linked immunosorbent assay (ELISA) kits for IL-8/CXCL8 (dilution 1:5; R&D Cat. # DY208-05, Lot #P105639) and CCL2/MCP-1 (dilution 1:15; R&D Cat. # DY279-05, Lot #333900) following the manufacturer’s instructions. Human Luminex analysis was performed using Luminex200 (SN LX10009028406) with a xPONENT 4.3 Software version with the following parameters: DD gate 8,000-16,500, 50 μl of sample volume, 50-100 events per bead/region, and Low PMT (LMX100/200: Default). All cytokines and chemokines analyses were measured at 24 h and 120 h post-infection. Some cytokines/chemokines levels were out of range, mainly reporting lower values. The dashed line (LLOQ and HLOQ) in those graphs indicates that the analyte levels are not necessary quantitatively determined with suitable accuracy.

### Statistical Analysis

An unpaired, 2-tailed Student’s *t*-Test for two-group comparisons was determined to assess statistical significance using GraphPad Prism 8 Software. In these studies, “n” values represent the number biological replicas using a different ALF sample from different adult or elderly human donors. Statistical differences between groups were reported as significant (*) when the p-value is less than or equal to 0.05.

## ACKNOWLEDGEMENTS

This study was supported by the National Institute on Aging (NIA), National Institutes of Health (NIH) (Grant number P01 AG-051428 to J.T., J.B.T., B.I.R and L.S.S.), and J.B.T. was partially supported by Robert J. Kleberg, Jr. and Helen C. Kleberg Foundation. The research reported in this publication was also partially supported by the Office of the Director, NIH under Award Number S10OD028653. The content is solely the responsibility of the authors and does not necessarily represent the official views of the National Institutes of Health. A.M.O.-F. was supported by the Douglass Graduate Fellowship at Texas Biomedical Research Institute.

## AUTHOR CONTRIBUTIONS

A.M.O.-F. and J.B.T. contributed to the design of the studies. A.M.O.-F., J.M.S. and A.G.-V. contributed to the experimental procedures. A.M.O.-F. and J.M.S. contributed to the acquisition of data and data analyses. J.I.P. and D.J.M. performed bronchoalveolar lavages (BAL) and provided the BAL fluid samples. D.L.C. provided TB-transmission electron microscopy expertise and scoring the samples by blinded analysis. J.B.T., L.S.S., J.T., D.L.C. and B.I.R provided a critical analysis of the data and editing of manuscript. A.M.O.-F. and J.B.T. wrote the manuscript.

## ADDITIONAL INFORMATION

### Supplementary information

The online version contains supplementary material available at

