## Supplementary Information for "Human alveolar lining fluid from the elderly promotes *Mycobacterium tuberculosis* growth in alveolar epithelial cells and bacterial translocation into the cytosol"

**Running Title:** Human lung mucosa influences *M.tb* growth in ATs.

**Keywords:** *Mycobacterium tuberculosis*; aging; alveolar epithelial cells; alveolar lining fluid; cytosol; tuberculosis.

### SUPPLEMENTARY INFORMATION

### SUPPLEMENTARY MATERIALS AND METHODS

***Ethics Statement and Human Subjects -*** Human donors with the following comorbidities were excluded: smokers (current or less than one month), excessive alcohol users, non-injection/injection recreational drug use, active pneumonia, asthma, bilateral cancer and on chemotherapy, COPD, pre-diabetes/diabetes (hemoglobin A1c higher than 5.7% within the last three months or a fasting blood glucose level of higher than 110 mg/dL), BMI equal or higher than 40, hepatitis, human immunodeficiency HIV/AIDS, immunosuppression or taking nonsteroidal anti-inflammatory agents (TNF antagonists), leukemia/lymphoma, liver failure, renal failure, rheumatoid arthritis, pregnancy (or gave birth less than three months ago), taken antibiotics recently and/or regular treatment with over the counter anti-inflammatory medications. Each “n” value represents a different human BALF donor (ALF), as specified in the figure legends.

***AT cell viability assay -*** At indicated times post-infection, AT cytotoxicity was determined by CellTiter-Glo® luminescent cell viability assay (Promega Cat. #G7570) following the manufacturer’s instructions. The assay consists of determining the number of viable cells in culture based on the quantification of ATP present in the media (the amount of ATP is directly proportional to the number of viable cells present in the culture media). The luminescent signal generated by the thermostable luciferase was measured every 24 h for up to 120 h with a GloMax Multi Detection System.

***AT compartment acidification assay -*** AT compartment acidification was determined by LysoTracker-Red® assay (Invitrogen Cat. #L7528), at 72 hours post-infection following the manufacturer's instructions. The assay consists of incubating the infected cells with red fluorescent acidotropic probes for labeling and tracking acidic organelles in live cells. Cells were visualized by laser scanning confocal microscopy using ZEISS LSM 800 microscope set at appropriate parameters and a final magnification of 600X. Quantification of co-localization of GFP-*M.tb* and LysoTracker-Red was determined by counting at least 100 events per condition in duplicate. All microscopy data were analyzed with Zeiss ZEN Software.

***Transmission electron microscopy -*** Infected ATs, at 72 hours post-infection were fixed in 2.5% glutaraldehyde and 2% formaldehyde (in 0.1 M Na Cacodylate pH 7.3) and analyzed by transmission electron microscopy (TEM). Briefly, after the primary fixative (overnight), the bacteria were rinsed with 0.1M phosphate buffer and post-fixed with 1% Zetterqvist's buffered Osmium Tetroxide for 30 min. Then, a stepwise prolonged dehydration procedure with a graded series of alcohols, all steps being for 10 min only one time, except 100% alcohol steps, which were 10 min twice, and finally dehydrated with propylene oxide twice for 10 min each. For resin infiltration and embedding, the cell blocks went from Propylene oxide to 3:1 Propylene oxide:Epon for 4 h incubation followed by two more incubations in 1:1 Propylene oxide:Epon and 2:1 Propylene oxide:Epon on a rotator. Cell blocks were transferred to neat Epon for 4 h and curing at 60°C for 48 h. Ultrathin sections were cut and post-stained on grid with 1% uranyl acetate in 50% MeOH for 1 h at room temperature in the dark followed by staining with lead citrate for 3 min at room temperature.

### SUPPLEMENTARY FIGURE LEGENDS

**Supplementary Figure S1. Differences in intracellular growth were not due to a significant decrease in cellular viability.** CellTiter-Glo® luminescent cell viability assay determines the number of viable cells in culture based on quantification of ATP present in the supernatants (Promega Cat. #G7570). The amount of ATP is directly proportional to the number of viable cells present in the culture. Representative experiment in triplicate of n=2 [Mean ± SEM], using two different A-ALFs and E-ALFs. Student's unpaired *t*-Test; Adult vs Elderly, *p<0.05, **p<0.01, ***p<0.001, ns: no significant differences. A: Adult exposed *M.tb*, E: Elderly exposed *M.tb.*

**Supplementary Figure S2. Effect of A- and E-ALF-*M.tb* on AT surface expression determined by mean fluorescence intensity (MFI).** Surface expression of infected ATs with either A-ALF or E-ALF exposed-*M.tb* were measured by flow cytometry. (**A**) Mean fluorescence intensity (MFI) relative fold change (versus uninfected ATs) of HLA-ABC (MHC Class I) expression overtime. (**B**) Mean fluorescence intensity (MFI) relative fold change (versus uninfected ATs) of HLA-DR/DP/DQ (MHC Class II) expression overtime. (**C**) No differences in bacteria inoculum used for the flow cytometry studies (percentage of cell expression and MFI analysis). Data shown are n=3 [Mean ± SEM], using three different A-ALFs and E-ALFs. Student's unpaired *t*-Test; Adult vs Elderly, *p<0.05, **p<0.01, ***p<0.001, ns (or absence of line): no significant differences. A: Adult exposed *M.tb*, E: Elderly exposed *M.tb*, UI: Uninfected ATs.

SUPPLEMENTAL FIGURES

FIGURE S1

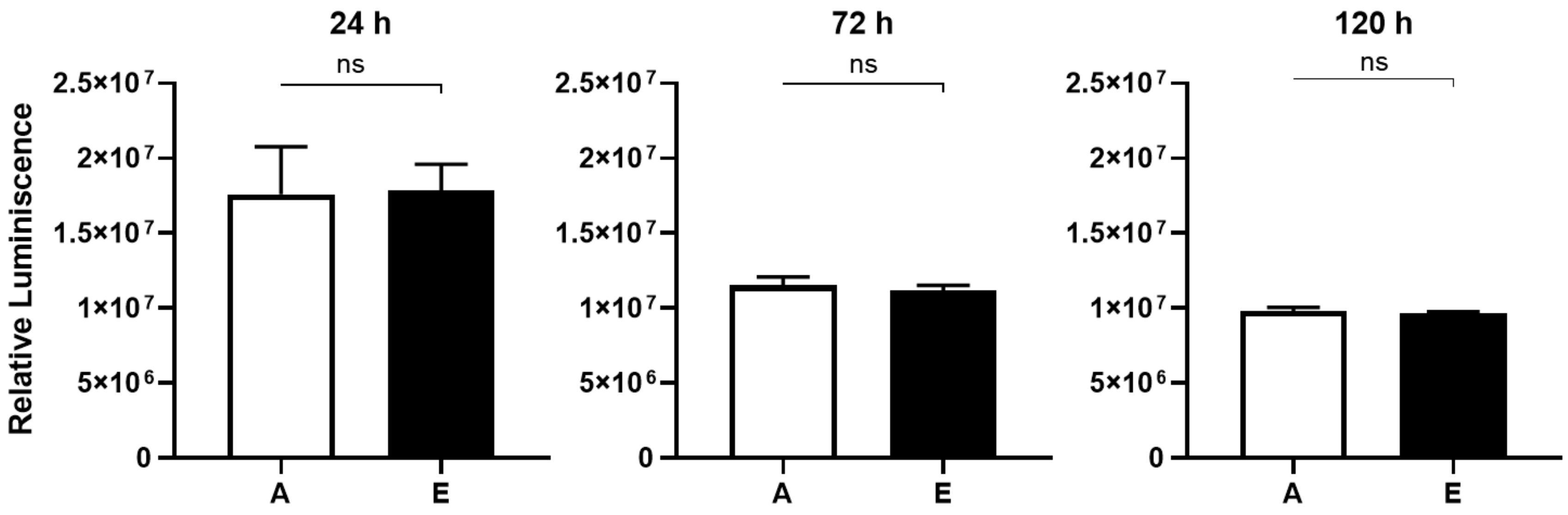

**FIGURE S2**

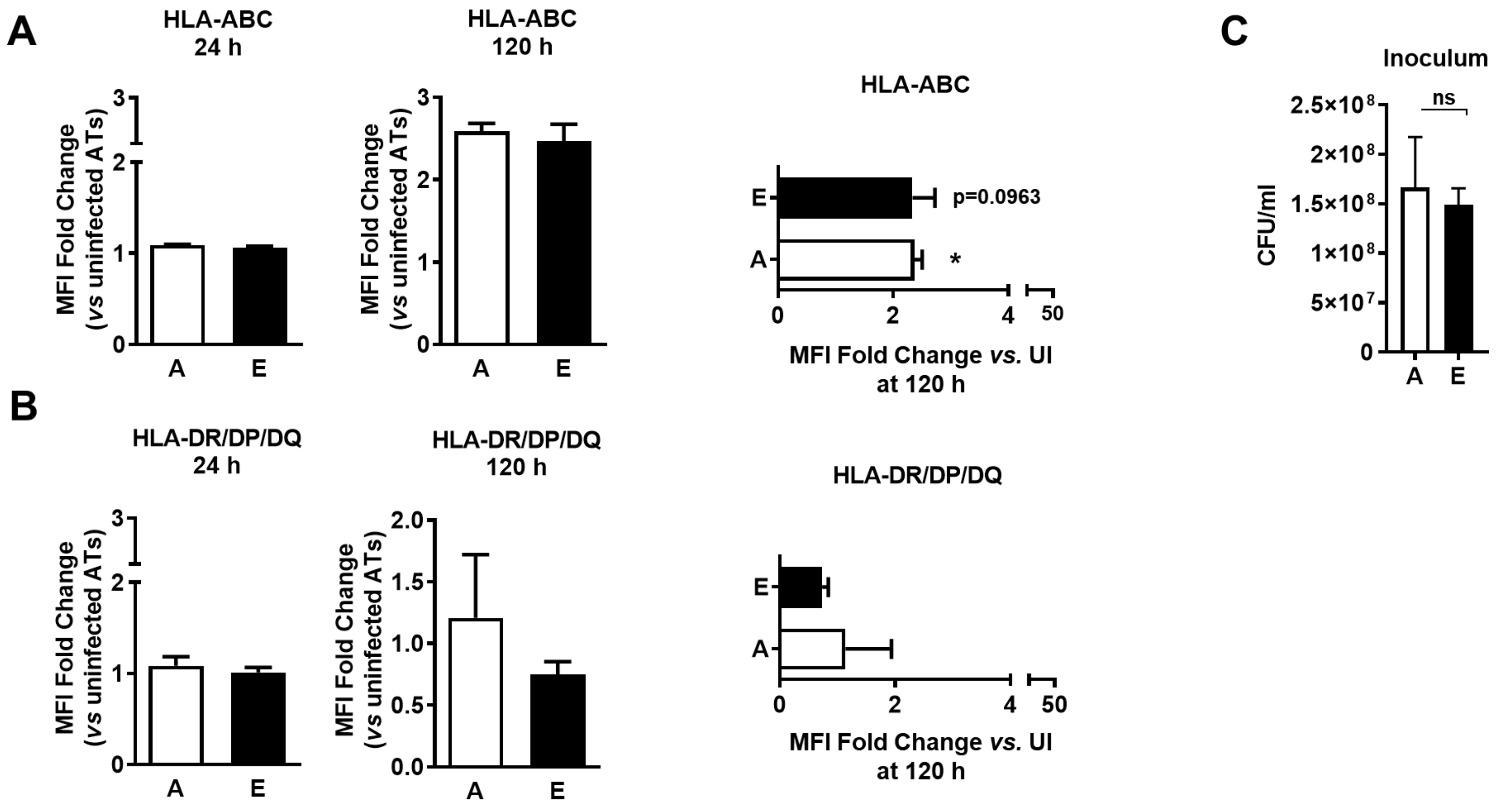

A
HLA-ABC
24 h
MFI Fold Change
(vs uninfected ATs)
3
2
1
0
A
E
HLA-ABC
120 h
MFI Fold Change
(vs uninfected ATs)
3
2
1
0
A
E
HLA-ABC
E
A
p=0.0963
*
0
2
4
50
MFI Fold Change vs. UI
at 120 h
B
HLA-DR/DP/DQ
24 h
MFI Fold Change
(vs uninfected ATs)
3
2
1
0
A
E
HLA-DR/DP/DQ
120 h
MFI Fold Change
(vs uninfected ATs)
2.0
1.5
1.0
0.5
0.0
A
E
HLA-DR/DP/DQ
E
A
0
2
4
50
MFI Fold Change vs. UI
at 120 h
C
Inoculum
CFU/ml
2.5×10^8
2×10^8
1.5×10^8
1×10^8
5×10^7
0
ns
A
E
